# CD14-deficiency protects against osteoarthritic subchondral bone sclerosis via enhanced osteoclastogenesis following joint injury

**DOI:** 10.64898/2026.02.10.705094

**Authors:** Lance A Murphy, Kate L Sharp, Kevin G Burt, Baofeng Hu, Vu Nguyen, Aurea R Borges, Christine Chung, Jonathan J Miner, Robert L Mauck, Carla R Scanzello

**Affiliations:** Perelman School of Medicine, University of Pennsylvania; Philadelphia, 19104, PA; Cartilage Regeneration using Advanced Technologies to Enable (CReATE) Motion Center, 10 CMC VA Medical Center; Philadelphia, 19104, PA; Department of Orthopaedic Surgery, University of Pennsylvania; Philadelphia, 19104, PA; Veterans Affairs Medical Center, San Diego, California; Department of Radiology, University of California San Diego, California; Division of Rheumatology, Perelman School of Medicine, University of Pennsylvania; Philadelphia, 19104, PA

**Keywords:** Type I Interferons, Osteoclast, Osteoblast, CD14, Toll-like Receptor 4, Osteoarthritis, Subchondral Sclerosis, Bone Remodeling

## Abstract

Aberrant bone remodeling is a hallmark of osteoarthritis, the most common arthritis affecting over 27 million US adults. Subchondral bone sclerosis, one sign of aberrant bone remodeling observable by routine x-rays, occurs as the trabeculae thicken, leading to increased bone volume. Toll-like receptors, pattern-recognition receptors of the innate immune system, have been implicated in OA pathogenesis, with TLR ligands, receptors, and co-receptors shown to mediate the severity and progression of OA. We have previously shown that CD14-deficiency protects mice against post-traumatic OA, and specifically reduces subchondral sclerosis post-injury. *We hypothesized that depletion of CD14 protects against TLR4-dependent inhibition of osteoclastogenesis and therefore increases OC density in the SCB after injury, mitigating aberrant bone deposition in a murine model of OA*. To determine how cellular changes correlate with bone structure derangements post-DMM, we performed MicroCT, Tartrate-resistant acid phosphatase staining, and alkaline phosphatase staining. To establish mechanistic changes in cellular signaling, we isolated WT and CD14-deficient osteoclast precursors and subjected them to LPS, an osteoarthritis-relevant TLR ligand, during differentiation. CD14-deficient mice, as well as WT mice treated with an anti-CD14 monoclonal antibody, show protection from post-injury increases in both bone volume fraction and bone mineral density. CD14-deficient mice had an increased osteoclast presence in the SCB two weeks post-injury, potentially protecting them from increases in bone volume and density. *In vitro*, CD14-deficient OCPs differentiated faster than WT OCPs, due to reduced Type I Interferon (IFN-I) signaling. In the presence of an LPS challenge, CD14-deficient OCPs were protected against LPS and TLR4-mediated inhibition, likely due to decreased MyD88-dependent TLR4 signaling. This work opens up new potential pathways to therapeutically target aberrant bone remodeling in the setting of joint injury and PTOA.

**Lay Summary:** Osteoarthritis is one of the leading causes of disability worldwide. One of the hallmarks is subchondral sclerosis, or thickening of the bone in and around the joint. In this work, we used a mouse model of osteoarthritis to show that decreasing inflammatory signaling, through removal of CD14, protects against subchondral sclerosis, due to an increased presence of osteoclasts, cells that combat bone thickening. Osteoclasts without CD14 differentiate faster than osteoclasts with CD14, due to decreased Type I Interferon, an inflammatory cytokine.

**Graphical Abstract:** 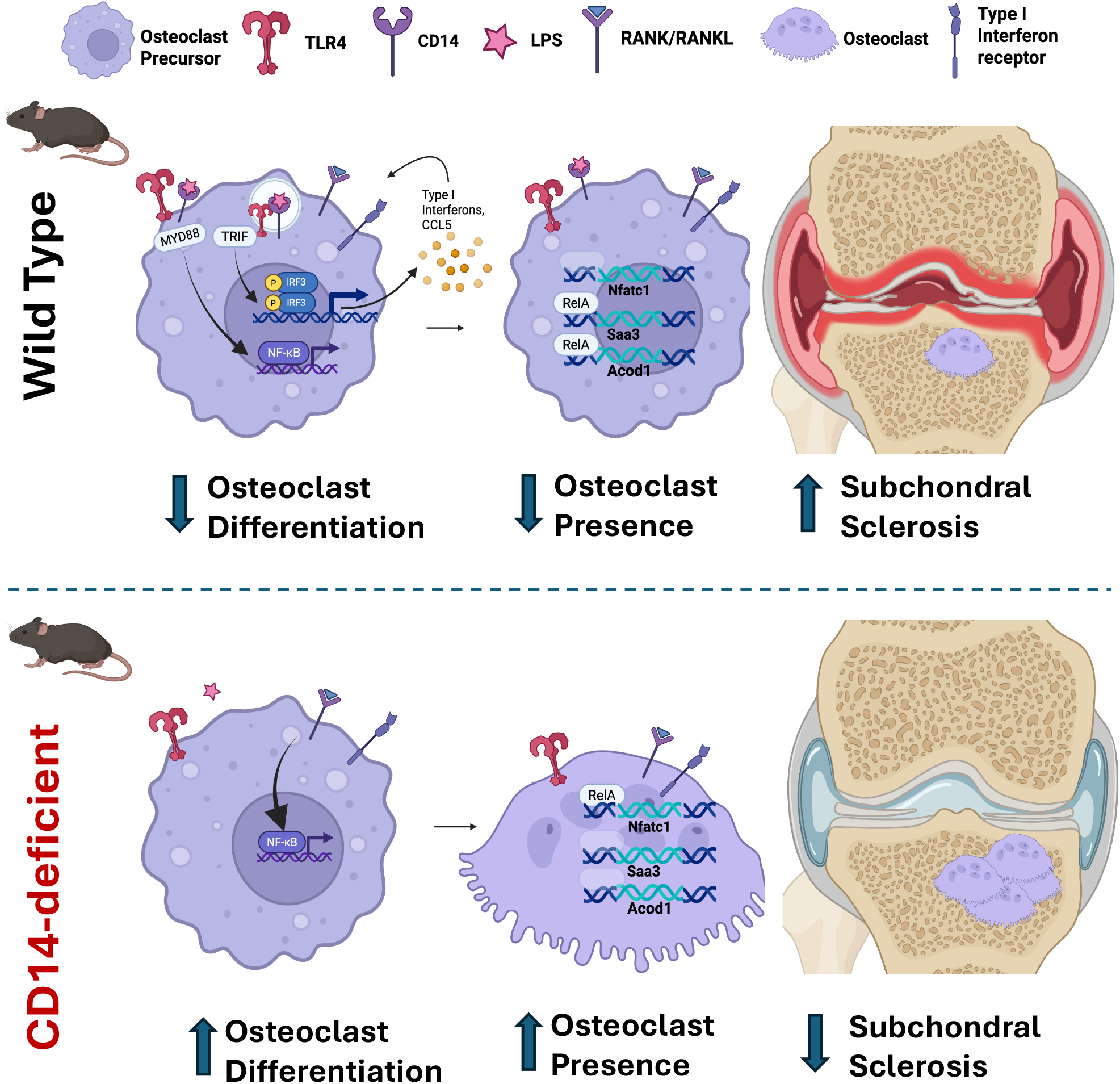

## Introduction

Aberrant bone remodeling is a hallmark of osteoarthritis (OA), the most common arthritis affecting over 27 million US adults. Signs of pathologic bone remodeling in OA include subchondral bone sclerosis, marginal osteophytes, and bone marrow lesions in the subchondral areas, all of which are associated with pain in patients.^1^ Subchondral sclerosis, observable by routine x-rays, occurs as the trabeculae thicken, leading to increased bone volume. However, these thickened trabeculae have poor structural quality as they are improperly mineralized.^1^ Osteophytes form through a process similar to endochondral ossification, beginning as chondrophytes which then hypertrophy, mineralize and ossify, leading to bone deposition at the margins of articular joints. Bone marrow lesions (BMLs), seen by magnetic resonance imaging (MRI), are characterized by complex histopathologic changes, including fibrosis, thickened trabeculae, and new cartilage formation.^2^ BMLs are additionally characterized by increased neuronal infiltration, hypervascularization, and inflammation, with a corresponding increase in osteoclasts which exhibit upregulated neurogenic, angiogenic, and inflammatory transcriptional activity.^3^ Aberrant subchondral bone (SCB) remodeling is often an early manifestation of OA in preclinical models ^4^ and can influence the health of the overlying articular cartilage via altered mechanical loading and direct bone-cartilage crosstalk.^5,6^ Thus, targeting bone remodeling could have implications for treating *both* structural pathology and pain in OA.

The innate immune system plays a major role in mediating bone remodeling in both homeostasis and disease.^7^ Toll-like receptors (TLRs) are pattern-recognition receptors of the innate immune system that respond to damage-associated molecular patterns (DAMPs) and pathogen-associated molecular patterns (PAMPs), leading to pro-inflammatory signaling.^8^ TLR pathways have been implicated in OA pathogenesis, with TLR ligands, receptors, and co-receptors implicated in the severity and progression of OA.^9^ For example, LPS (a TLR4 ligand) is present at higher levels in the synovial fluid and in the systemic circulation in patients with OA.^10^ Furthermore, soluble TLR4 in plasma is associated with the progression of cartilage damage over 18-months.^11^ Synovial fluid and plasma content of CD14, which functions as a cell surface and soluble co-receptor for several TLRs, is associated with pain as well as presence and progression of osteophytosis in patients suffering from OA.^12^ CD14 potentiates and sensitizes the response to TLR ligands and can mediate cartilage destruction in the destabilization of the medial meniscus (DMM) murine model of OA.^13,14^ We previously demonstrated that CD14-deficient mice were protected from progressive cartilage damage and decreased mobility in the DMM model.^15^ Interestingly, CD14-deficiency also significantly protected mice against pathologic subchondral bone deposition after DMM injury.

To understand the mechanism by which CD14-deficiency protects against bone remodeling, it is important to understand its effects on osteoclasts (OCs) and osteoblasts (OBs). Prior literature suggests that TLR4 activation by LPS has a divergent effect on osteoclastogenesis, depending on the status of the cells in their differentiation spectrum.^16,17^ For example, TLR4 activation in cells already primed with RANKL is pro-osteoclastogenic. Conversely, TLR4 activation in OC precursors that are not RANKL-primed is inhibitory to osteoclastogenesis. Although TLRs and CD14 are most highly expressed on cells of myeloid lineage, including osteoclast precursors (OCPs), there are also implications for mesenchymal lineage cells that mediate bone remodeling, specifically osteoblasts.^14^ At homeostasis, osteoblastic TLR4 signaling is necessary for loading-induced bone formation in mice.^18^ Moreover, TLR4 signaling in OBs leads to the production of RANKL, which in turn stimulates osteoclastogenesis.^19^

Given this complex role, we sought to understand the cellular activities and signaling that underlies the differential bone remodeling observed after joint injury in mice deficient in CD14. To do so, we evaluated how CD14-deficiency impacts OC and OB density in the SCB after joint injury, and the impact of CD14-deficiency on OC differentiation and activation *in vitro. We hypothesized that depletion of CD14 would protect against TLR4-dependent inhibition of osteoclastogenesis and therefore increase OC density in the SCB after injury, mitigating aberrant bone deposition in a murine model of OA*.

## Materials and Methods

### Genetic ablation of CD14 in the context of joint injury

Male C57BL/6 (WT) and CD14KO mice (n=4-5 per group) underwent DMM or sham surgery at 12 weeks of age. Knees were harvested at 2, 4, and 8 weeks post-surgery and fixed in 4% PFA for micro-CT imaging. Knees were then embedded in OCT and cryosectioned using the tape stabilization technique for quantification of OC and OB content, as described below.^20^

### Pharmacologic inhibition of CD14 in the context of joint injury

In another related study, 28 male WT mice underwent DMM surgery, followed by intra-articular treatment with either a neutralizing monoclonal antibody (mAb) against murine CD14 (0.5 mg/kg, clone big53, provided by Implicit Biosciences, Ltd) or an IgG2a isotype control (n=14 per group). Mice received 3 weekly doses starting 48-hours post-DMM surgery. After 12 weeks, mice were sacrificed and knee joints were fixed in 4% PFA and imaged by micro-CT.

### Micro-CT evaluation of SCB structure

#### CD14 genetic deficiency study

*Ex vivo* bone assessments of the knee were performed using micro-computed tomography. At 2, 4, and 8 weeks, analysis was completed as previously described.^15^ Briefly, dissected knees (n = 4–6) were scanned using a VivaCT -40 micro-CT scanner (Scanco-Medical, Brüttisellen, Switzerland) at a voxel size of 5 μm, energy of 55kV, intensity of 109 μA, and an integration time of 300 ms. The medial epiphysis of the proximal tibia served as the region of interest given that changes in this region have been reported at the timepoints used in this study.^21^ The 3-dimensional reconstructed region was manually contoured to include calcified cartilage and extend to the growth plate. To avoid osteophytes, ten percent of the medial and lateral width of the tibial plateau was excluded. Manual segmentation was repeated every tenth slice and splined together automatically to create a volume of interest (VOI). Percent bone volume per total volume (% BV/TV), bone mineral density (BMD, mg HA/cm^3^) and trabecular thickness (Tb.Th., mm) was calculated using the manufacturer’s (ScanCo, Inc.) software.

#### CD14 pharmacologic blockade study

Micro-CT scans (SkyScan 1174, Bruker) from a separate study examining the effects of pharmacologic blockade on post-DMM joint pathology (PMID 40502022) were assessed for SCB remodeling at 12 weeks post-injury. Scanning parameters included a voxel size of 9 um, an energy of 50 kV, an intensity of 793 µA, and an integration time of 1600 ms. A calcium hydroxyapatite (HA) phantom with two reference concentrations (0.25 and 0.75 g HA/cm^3^) was scanned alongside each sample for mineral density calibration. Bone analysis was limited to the femur due to technical challenges with tibial subchondral bone segmentation and quantification. Analysis was performed using a custom MATLAB script. Bone mineral density (BMD, g/cm^3^) and bone volume fraction (BV/TV, %), were quantified in regions of interest (ROIs) selected in the sagittal plane to capture the majority of each femoral condyle, with emphasis on the central region. Manual segmentation was performed to define the subchondral bone.

### Tartrate Resistant Acid Phosphatase (TRAP) and Alkaline Phosphatase staining

Knee sections from WT and CD14-deficient mice were stained with ELF97 (ThermoFisher) phosphatase substrate under acidic conditions (to identify TRAP+ OCs), or basic conditions (to identify Alk Phos+ OBs).^22,23^ TO-PRO-3 Iodide was used as a nuclear counterstain. Sections were imaged using an Axioscan 7 slide scanner (Zeiss, 20X objective). The medial tibial plateau was chosen as the region of interest, and OC and OB surface lining distances were measured and normalized to the total trabecular surface using the freehand line and measurement features in ImageJ (v1.54).

### *In vitro* OC differentiation assay

Bone marrow was collected from the femorae and tibiae of 12-week old male mice (WT/C57BL/6 and CD14KO) pooling cells from 3-5 mice per specimen. Additionally, Interferon Alpha Receptor 1-deficient (IFNAR1KO) and Interferon Regulatory Factors 3 and 7 double knockout (IRF3/7DKO) mice were obtained from a collaborator and cells were isolated from bone marrow as mentioned above. Cells were cultured in αMEM with 30 ng/mL M-CSF for 5-7 days to expand osteoclast precursors (OCPs). OCPs were then replated at 50,000 cells per well (24-well plates) in the presence of 30 ng/mL M-CSF and 100 ng/mL RANKL. Supernatants were removed for LPS quantification and cells were fixed on days 3 and 4 and stained using the Leukocyte Acid Phosphatase Kit (Sigma) for TRAP positivity. To quantify OC number, osteoclasts were considered to be any TRAP+ cell with greater than 2 nuclei. Three images were acquired per well. Osteoclasts were manually traced and the percentage area of each 10X field covered by OCs was quantified using CellProfiler. The percentage of area covered was then averaged for each well and normalized to WT average.

For select experiments, cells were fixed on days 3-6 after RANKL addition, stained with Rhodamine Phalloidin (for F-actin) and DAPI (for nuclei), and imaged by confocal microscopy (10x), as an additional method for quantifying OCs.^24^ The percentage of area covered by OCs was analyzed in the same manner as the TRAP staining.

### Osteoclast functional assay

Osteoclast functional activity was assessed using the Bone Resorption Assay Kit (CosmoBioUSA). After expansion, osteoclast precursors were plated onto a calcium phosphate-coated plate loaded with fluorescently-associated chondroitin sulfate (FACS) and exposed to M-CSF and RANKL as described previously. FACS released into the media was measured using a fluorescent plate reader (emission of 520 nm) as a measure of resorption activity, on days 3, 4, 5, and 6 post-RANKL stimulation. After lysis of cells, wells were stained with toluidine blue to visualize resorptive pits. Pits were imaged and pit area was quantified and expressed as the percentage of total area per well.

### Bulk RNA sequencing

RNA was harvested from WT and CD14-deficient OCs 4 days after the addition of RANKL using the Qiagen RNeasy kit, and submitted to the Penn Genomics and Sequencing Core. After sequencing, quality control analyses were completed with Fastqc. Exploratory pathway analysis was completed by mapping gene sets to the Hallmark pathway set from Molecular Signatures Database.

### Effects of LPS, TLR4, and Type-I IFN on osteoclastogenesis

OCPs were treated with LPS, CLI-095 (1 µg/mL, TLR4 inhibitor), and/or anti-IFNAR1 (1 µg/mL) antibody during RANKL exposure. After 4 days, cells were stained for TRAP and imaged and quantified as above. In additional experiments, RNA was isolated from WT and CD14-deficient osteoclasts on day 4 for PCR analysis.

### RT-qPCR

RNA from WT and CD14-deficient osteoclasts was isolated on day 4 post-RANKL stimulation using the Qiagen RNeasy Mini Kit according to the user manual and cDNA was generated using the Bio-Rad iScript CDNA Synthesis Kit. Real-time quantitative PCR was then completed using iQ SYBR Green Supermix (Bio-Rad) and the following primers: *Saa3 (*Fwd: 5’-GCC TTC CAT TGC CAT CAT TC-3’ Rev: 5’-CAT ATG TCT CTA GAC CCT TGA C-3’), *Acod1 (*Fwd: 5’-GGC GTT CAA CGT TGG TAT TG-3’ Rev: 5’-AGG GTG GAA TCT CTT TGG TAT G-3’), *PRDX1 (*Fwd: 5’-TTT ACC TGC CTG TTG GAT ACC-3’ Rev: 5’-GGA GAG ACA GCT CAA TGG TTA G-3’), *NFATc1 (Fwd: 5’-CCT CTG TGA GTC TTT GGG TTA G-3’ Rev: 5’-ACC ACG GCA GGC TTA TTT-3’)*, and *TBP (Fwd: 5’-CTA CCG TGA ATC TTG GCT GTA A-3’ Rev: 5’-GTT GTC CGT GGC TCT CTT ATT-3’)* used as the reference gene.

### Statistics

One-way ANOVA was used with unpaired t-tests and Holm-Sidak correction for pairwise comparisons. For studies comparing effects of both mouse strain and treatment, a two-way ANOVA was used. All statistical analyses were performed using GraphPad Prism 10.0 (GraphPad Software Inc., La Jolla, CA, United States). Results were expressed as mean ± SD.

## Results

### SCB remodeling after joint injury in the context of CD14-deficiency or blockade

In order to understand the effect of CD14-deficiency on SCB remodeling, we first evaluated time-dependent changes in SCB volume, evaluated by microCT, in the tibial subchondral region for up to 8 weeks following DMM injury, comparing WT and CD14-deficient strains. As shown in **Figure 1A**, while WT mice showed a 29% increase in total bone volume fraction (BV/TV) by 2 weeks post-DMM (p<0.01), CD14-deficient mice showed a delay in this increase until 4 weeks post-DMM, with observable changes on microCT images (**Fig. 1B**). At the 2-week timepoint, WT DMM-operated mice showed a 100% increase in trabecular thickness (p < 0.001) and a 13% increase in bone mineral density (p<0.01) compared to naïve and sham operated controls, while no increase in these parameters was seen after DMM-injury in the CD14-deficient strain (**Fig. 1C**). Compared to WT DMM mice, CD14-deficient mice that underwent DMM showed a 19% decrease in bone volume fraction (p<0.01), a 56% decrease in trabecular thickness (p<0.0001), and a 12% decrease in bone mineral density (p<0.01). These data suggest that genetic ablation of CD14 limits pathologic bone remodeling following joint injury.

**Figure 1:**
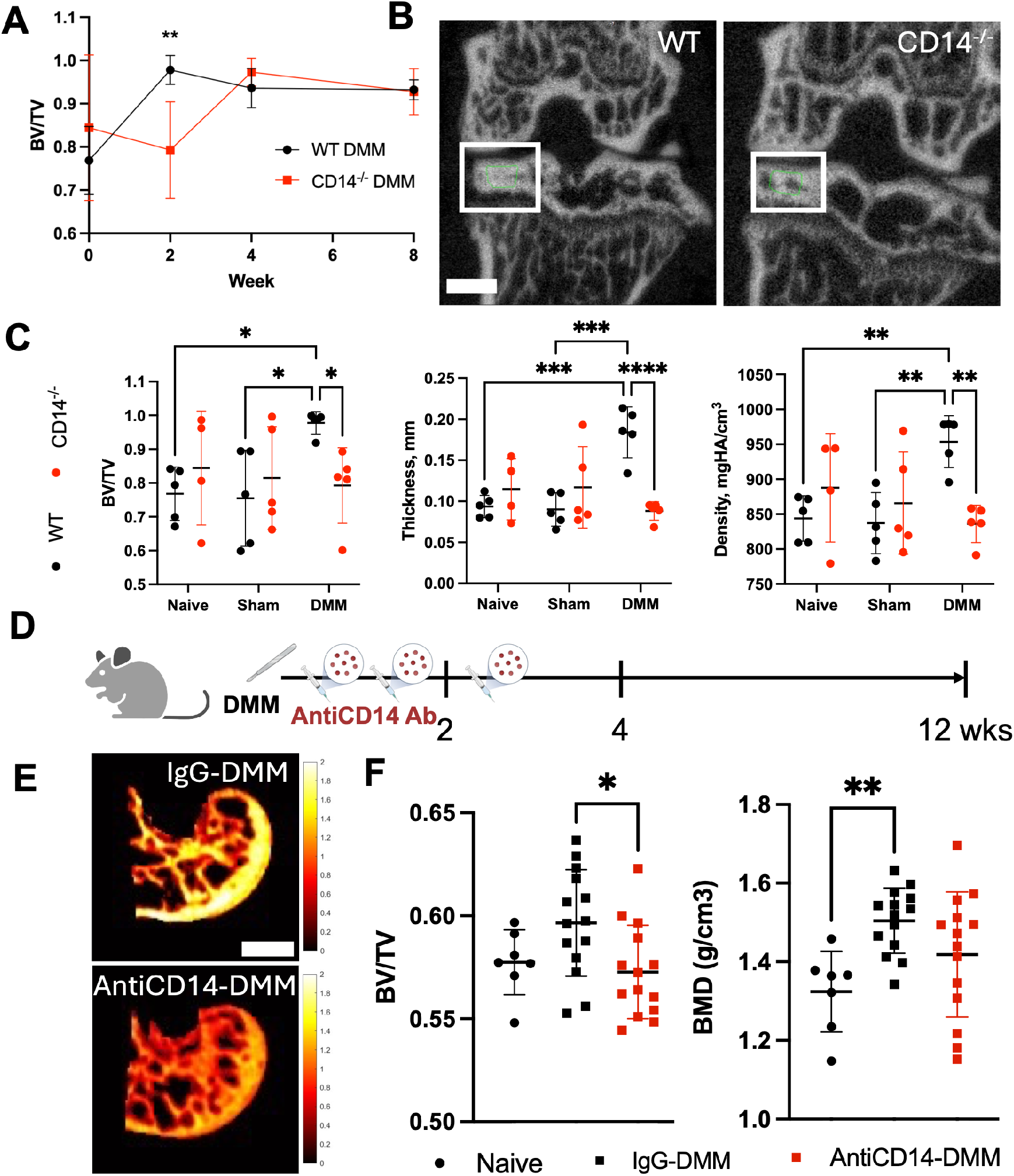
CD14-deficiency and pharmacologic blockade delays pathologic subchondral bone remodeling after int injury. Bone volume fraction (A, BV/TV) as a function of genotype and time post-DMM. Representative images of T (B, left) and CD14KO (B, right) DMM microCT images at 2 weeks post-DMM. Scale bar = 500 µm. White box utlines the area of interest. Bone volume fraction (C, left), trabecular thickness (middle), and bone mineral density ght) for naïve and injured (Sham vs DMM) WT and CD14KO mice at 2 weeks post-DMM. (D) Experimental schematic r pharmacologic CD14 inhibition study. Representative images of posterior medial femoral condyle of untreated (top) nd treated (bottom) mice. Scale bar = 500 µm. Bone volume fraction (F, left) and bone mineral density (F, right) of MM mice (with and without CD14 inhibition) and naïve controls. *p<0.05, **p<0.01, ***p<0.001, ****p<0.0001

To explore the translational potential of this finding, we next evaluated microCT scans from a study testing pharmacologic blockade of CD14, focusing on longer-term outcomes (12 week endpoint). Mice underwent DMM and were treated with a neutralizing anti-CD14 mAb weekly three times starting 2 days post-injury, or with an isotype-matched control mAb (**Fig. 1D**).

Focusing now on the femoral subchondral bone, which has been shown to exhibit changes only at later timepoints like 12 weeks post-injury, DMM-operated, control-treated mice showed significantly increased bone mineral density (13% increase, p <0.01) and slightly increased bone volume fraction from 57.7% to 59.7% (p= 0.1855) compared to age-matched naïve mice.^21^ Mice treated with the anti-CD14 mAb were protected from the increased bone mineral density and showed significantly decreased bone volume fraction compared to control-treated mice (4% decrease, p < 0.05) at 12 weeks post-DMM (**Fig. 1E&F**).

### Effects of CD14-deficiency on SCB osteoclast and osteoblast density post-DMM

Given the significant differences in the time course of post-DMM tibial SCB remodeling between WT and CD14KO mice, we next assessed OC and OB density within the medial tibial SCB compartment. We observed no differences between strains in terms of OB density (measured by Alkaline Phosphatase staining) at the 2 week timepoint, although there was an increase in CD14-deficient mice at 8 weeks (**Fig. 2A&C**).

Conversely, TRAP staining revealed a four-fold increase in osteoclast density in CD14-deficient compared with WT mice at 2 weeks (p<0.001, **Fig. 2B&C**), when strain-related differences in microCT measured bone volume were most pronounced (**Fig. 1A**). The increased osteoclast presence led to a 70% decrease in the OB:OC ratio (p<0.05, **Fig. 2D**, right) in the CD14-deficient strain compared to WT.

**Figure 2:**
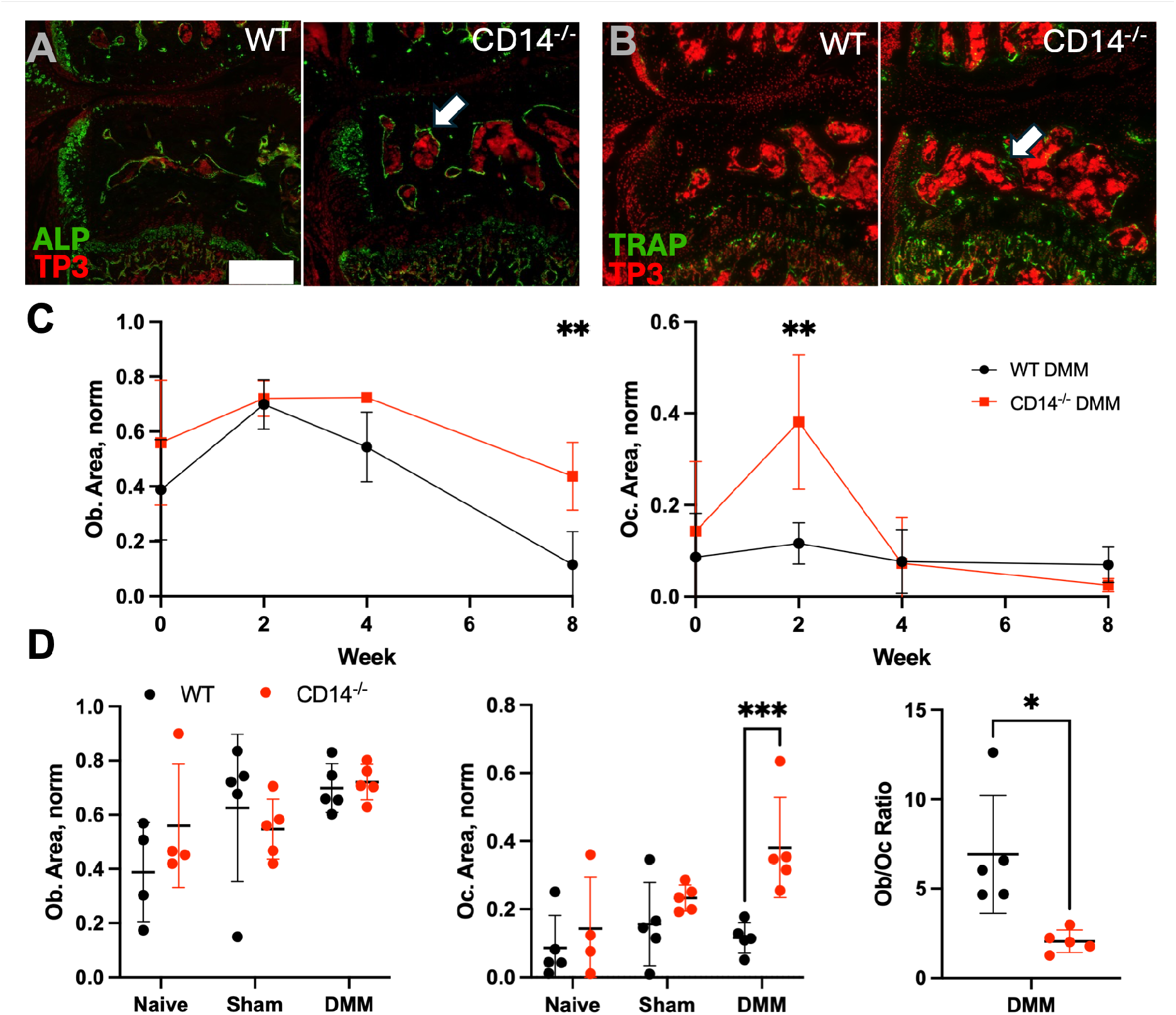
CD14-deficiency increases osteoclast presence in subchondral bone at 2 weeks post-DMM. Representative images of ALP staining in the medial tibial plateau of DMM WT (A, left) and DMM CD14KO (A, ght); arrows indicate regions of positive staining. Scale bar = 100 µm. Representative images of TRAP staining in the medial tibial plateau of DMM WT (B, left) and DMM CD14KO (B, right). Area covered by osteoblasts (C, left) and osteoclasts (C, right) in tibial SCB WT and CD14KO mice over 8 weeks post-DMM. Osteoblast area (D, left), osteoclast area (D, middle), and Ob/Oc ratio (D, right) for WT and CD14KO mice at wo weeks. *p<0.05, **p<0.01, ***p<0.001

### The effect of CD14-deficiency on RANKL-mediated osteoclast differentiation and function

As OCPs are known to express high levels of CD14, and the SCB cellular analysis indicated an increase in OC density in CD14-deficient mice, we next evaluated OC differentiation potential as a function of CD14. *In vitro*, CD14-deficient OCPs differentiated into OCs within 3 days of RANKL addition, while WT OCPs were delayed in their differentiation until day 4 (**Fig. 3A**). This was evidenced by an increased number of TRAP+ cells per well (14-fold, p<0.001), and increased percentage of the well area covered by TRAP+ multinucleated cells on day 3 (15-fold, p<0.001) and 4 (1.3-fold, p<0.05) in CD14KO vs. WT cells (**Fig. 3B)**. Osteoclasts are known to have a limited lifespan *in vitro*. To determine whether this lifespan was influenced by CD14, we extended the time course of analysis to 6 days following RANKL addition and measured OC differentiation via fluorescent Rhodamine staining of actin rings. As OCs mature, their actin rings become more pronounced, they grow larger, and they fuse to become multinucleated. Using this assay, CD14-deficient cells again reached peak OC differentiation at day 4, evidenced by mature actin rings, increased size and increased DAPI intensity, but then decreased at day 5, whereas WT controls reached peak differentiation at day 5 and decreased at day 6 (**Fig. 3C, D, and S1**). This suggests that the OC lifespan was similar between strains, despite CD14-deficient cells differentiating earlier *in vitro*. Next, the resorptive function of OCs over six days was measured, using a fluorescence-based assay. Although there was a trend towards an early increase in resorption on day 3 in the CD14-deficient OCs (1.2-fold, p =0.064), cumulative resorptive activity over six days was similar between strains (**Fig. 3E-G**). To establish whether the differences observed were a direct consequence of CD14 bioavailability, we added recombinant soluble CD14 to CD14-deficient cultures during the differentiation window. Addition of exogenous CD14 to CD14-deficient cultures inhibited osteoclast differentiation in a dose-dependent manner (**Fig. 3H**),reaching WT levels at the highest dose. Taken together, these data suggest accelerated OC differentiation in CD14-deficient cells, but no significant effect on OC survival or resorptive capacity.

**Figure 3:**
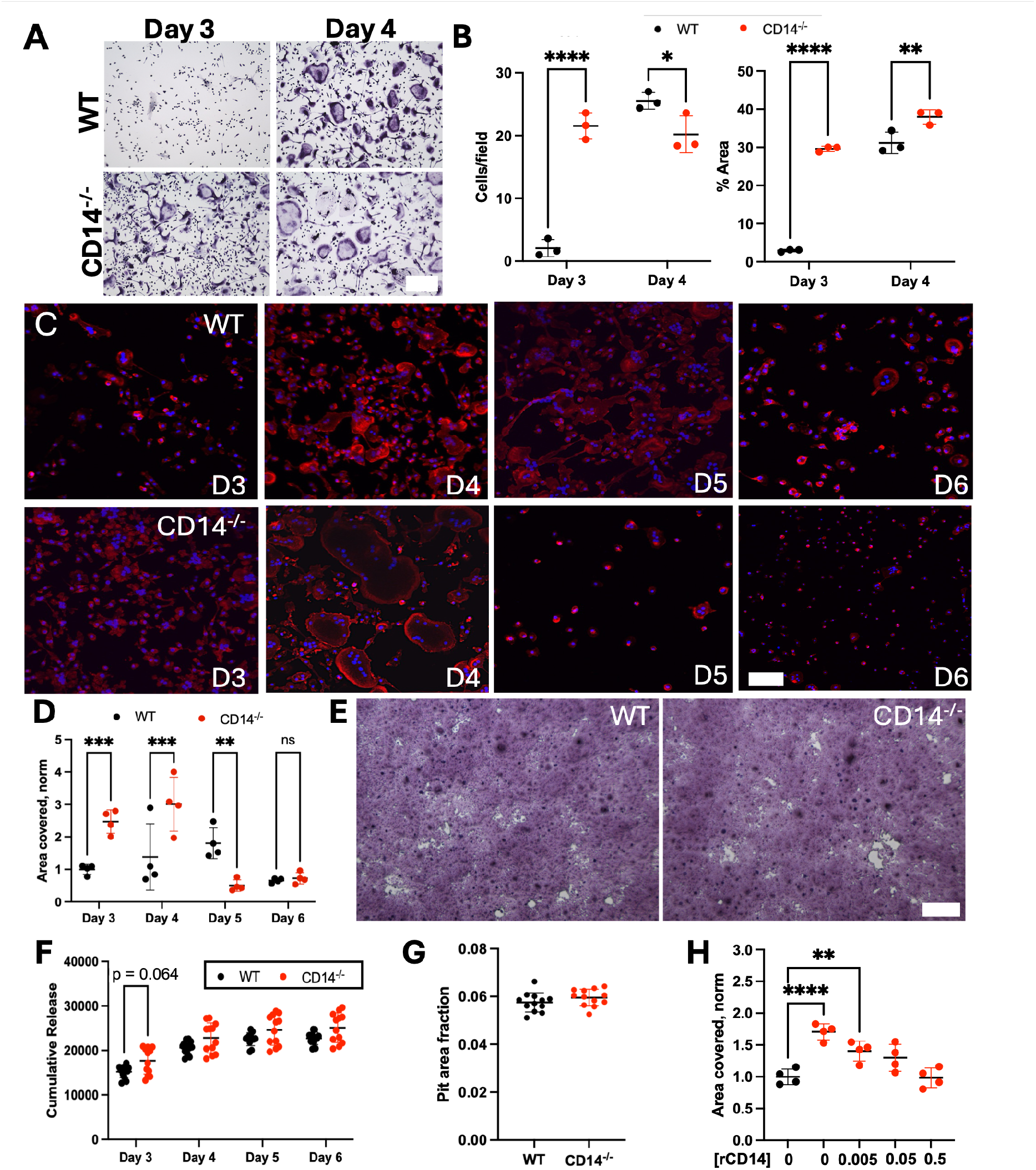
CD14-deficient osteoclasts differentiate faster in response to RANKL addition. Representative mages of WT and CD14KO osteoclasts on Day 3 and 4. Scale bar = 100µm (A). Quantification of osteoclast cells per field of view and % area covered (B). Representative images of WT and CD14KO osteoclast actin staining on ay 3, 4, 5, and 6 post-RANKL addition (C). Percent area covered by osteoclasts. Scale bar = 100 µm (D). epresentative images of resorption assay pits (E) for WT (left) and CD14KO (right) cultures. Scale bar = 500 µm. Cumulative release of fluorescent substrate (F) and pit area fraction on day 6 (G). Percent area of osteoclasts with the addition of recombinant CD14 (rCD14, H), from 0-0.5 μg/mL. *p<0.05, **p<0.01, ***p<0.001, ****p<0.0001

### CD14-deficient osteoclasts differentiate earlier due to decreased IFN-I signaling

In order to further understand the differences in OC differentiation, RNA from WT and CD14-deficient osteoclasts was isolated at day 4 post-RANKL stimulation and subjected to bulk RNA sequencing. Bulk sequencing revealed 2798 differentially expressed genes between strains (padj<0.01, **Fig. 4A**). Pathway analysis of this differential gene set revealed 11 pathways that were up-regulated and 10 that were down-regulated in CD14-deficient OC, including “MTORC signaling,” “Protein Secretion” and “Oxidative Phosphorylation”. Of particular interest was a decrease in Interferon signaling pathways (“Interferon Alpha Response” and “Interferon Gamma Response”), as it is well established that interferon signaling is inhibitory to osteoclast differentiation (**Fig. 4B&C**). Further evaluation of the differentially expressed genes (DEGs) within the two IFN pathway gene sets revealed 52.4% similarity in the genes within these pathways, including genes known to be inhibitory to OC differentiation (i.e. guanylate-binding protein 4 (GBP4) and GBP2) (**Fig. 4D**).^25^

**Figure 4:**
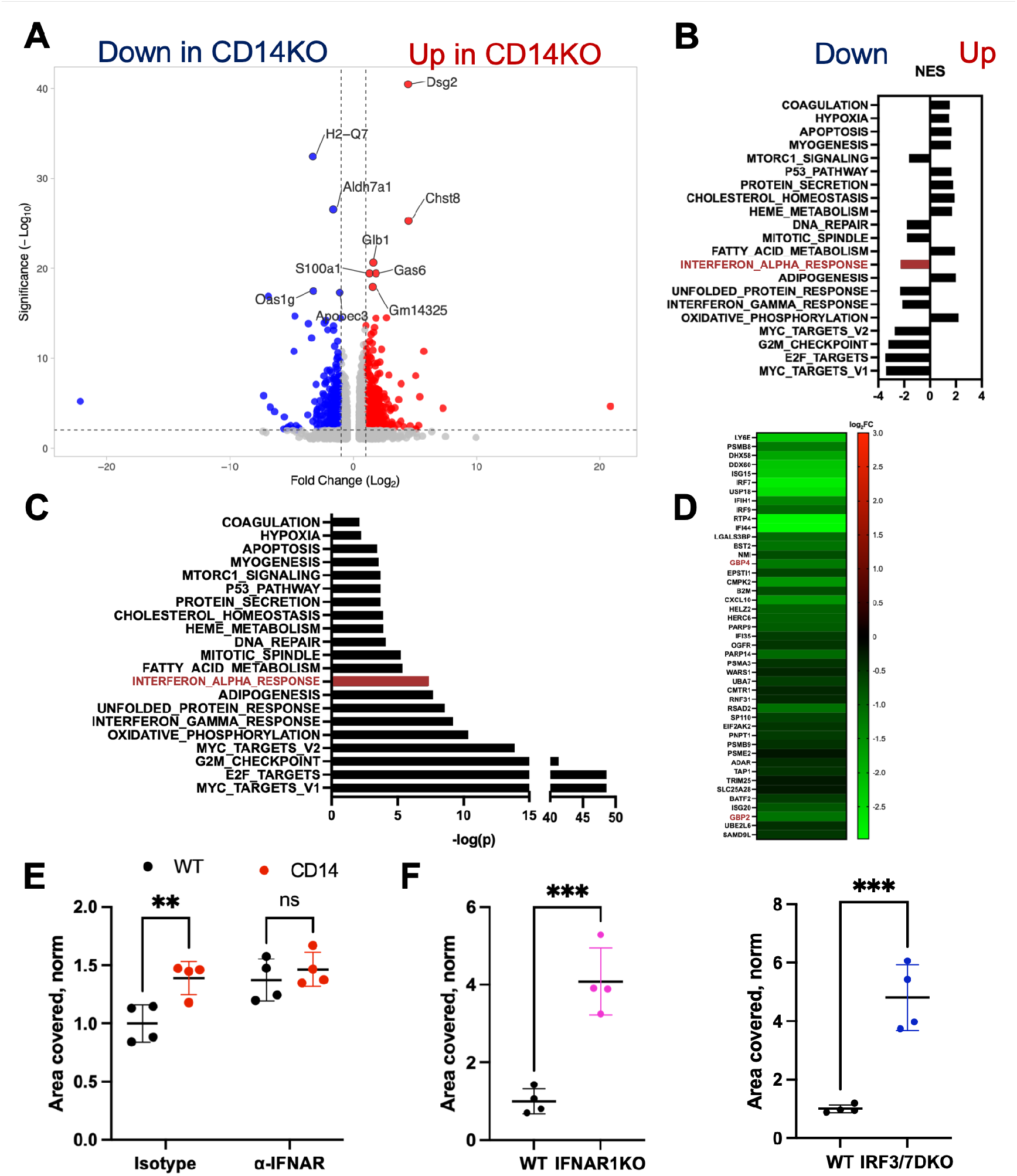
CD14-deficient osteoclasts differentiate earlier due to decreased IFN-I signaling. Volcano plot of differentially expressed genes (DEGs) between WT and CD14-deficient strains (A). Hallmark pathway analysis normalized enrichment score based on DEGs between strains (B), p-value (C), and leading edge genes for Interferon Alpha Response pathway (D). Percent area of WT and CD14KO osteoclasts with the dition of Anti-IFNAR1 antibody (E). Percent area of osteoclasts at day 3 after RANKL addition, for OCPs derived from WT, IFNAR1KO, and IRF3/7DKO. **p<0.01, ***p<0.001

Given the downregulation of IFN-I signaling in CD14-deficient osteoclasts, as well as the importance of CD14 in mediating IFN-I (including IFNβ) production downstream of TLR4, we hypothesized that the effects of CD14-deficiency on OC differentiation *in vitro* might be mediated by a reduction in IFN-I signaling.^14,26^ To test this, we treated WT cells with a neutralizing anti-murine IFNAR1 (receptor for IFN-I) antibody. This treatment increased WT OC differentiation to the level of CD14-deficient OC differentiation at day 3, while treatment with an isotype control did not (**Fig. 4E**). Treatment with the anti-IFNAR1 antibody did not affect CD14-deficient OC differentiation. Furthermore, osteoclasts derived from IFNAR1KO and IRF3/7DKO mice demonstrated increased differentiation capacity at day 3 compared to WT cells, phenocopying the effects of CD14-deficiency (**Fig. 4F**). These findings suggest that the accelerated rate of differentiation of CD14-deficient OCPs is due to a reduction in the production of IFN-I.

### Impact of CD14-deficiency on OC differentiation in the context of TLR ligand challenge

As CD14 is an important TLR4 cofactor and the osteoarthritic joint contains many potential TLR4 ligands, we next compared OC differentiation between WT and CD14-deficient cells in the presence of a TLR4 challenge. One TLR4 ligand thought to play a role in OA pathogenesis is the canonical TLR4 ligand, lipopolysaccharide (LPS).^27^ For physiologic relevance, we used LPS at a low concentration (1 ng/mL), consistent with concentrations observed in OA synovial fluid.^10^ Addition of LPS reduced OC differentiation in WT cells, as indicated by a 65% reduction in percent area covered at day 3 (p = 0.0558) and a 67% reduction at day 4 (p <0.01). LPS had no significant inhibitory effect on CD14-deficient cells (**Fig. 5B**). To confirm that the inhibitory effects of LPS were TLR4-mediated, we treated cells with a TLR4 inhibitor (CLI-095). This treatment partially rescued the inhibitory effects of LPS on WT OCs (**Fig. S2)**. In order to understand if these effects were also mediated by IFN-I, we again treated WT OCP with the anti-IFNAR1 antibody, this time in the presence of LPS. In this case, the anti-IFNAR1 antibody did not rescue WT OC differentiation (**Fig. 5C**). Furthermore, unlike CD14-deficient cells, OC differentiation in the IRF3/7KO and IFNAR1KO was still significantly inhibited by LPS treatment (**Fig. 5D & S3**). Together, this suggests that LPS-induced inhibition of OC differentiation in WT OCPs, and protection from LPS-induced inhibition in the CD14KO OCPs, is not mediated through IFN-I.

**Figure 5:**
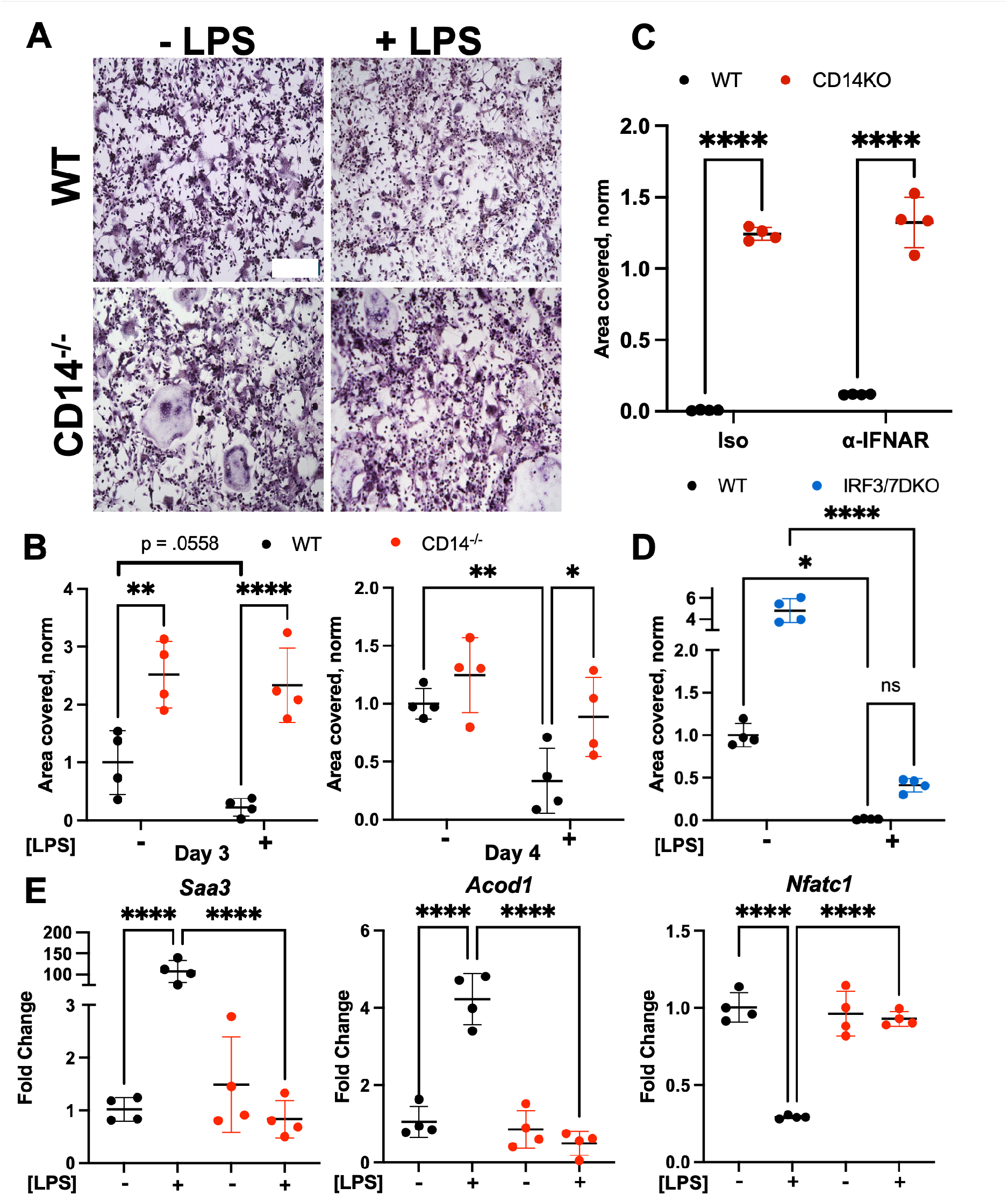
CD14-deficient osteoclasts are protected from LPS-induced inhibition. Representative images of T and CD14KO osteoclasts with and without LPS (A), with percent area covered by osteoclasts on day 3 and 4 ost-RANKL stimulation (B). Scale bar = 100 µm. Percent area of WT and CD14KO osteoclasts treated with LPS and Anti-IFNAR1 antibody (C). Percent area of WT and IRF3/7DKO osteoclasts with and without LPS (D). Transcript levels of *Saa3, Acod1*, and *Nfatc1* in WT and CD14KO osteoclasts with and without exposure to LPS). *p<0.05, **p<0.01, ****p<0.0001

Activation of TLR4 leads to cellular signaling cascades via two main pathways, a MyD88-dependent pathway which activates NF-kB driven transcriptional programs, and a MyD88-independent (and TRIF-dependent) pathway which leads to activation of both NF-kB and IRFs (Interferon regulatory factors).^28^ The MyD88-independent pathway is responsible for activation of the IFN-I response downstream of TLR4. Although it is known that IFN-I signaling is inhibitory to osteoclastogenesis, it has been reported that RANKL-mediated OC differentiation can be inhibited through both MyD88-dependent and -independent pathways downstream of TLR4 stimulation.^28,29^ In order to probe the effects of both TLR4-signaling pathways on OC differentiation, OCPs were isolated from TRIFKO and MyD88KO mice and cultured in the presence of RANKL. Both TRIFKO and MyD88KO precursors showed minimal capacity to differentiate into OC, suggesting a necessity for MyD88 and TRIF in OC differentiation *in vitro* (**Fig. S4)**.

Signaling through the MyD88-dependent pathway has been shown to inhibit OC differentiation by diverting NF-κB away from *Nfatc1* (Nuclear Factor of Activated T-cells, cytoplasmic 1, the master transcriptional regulator of OC differentiation), and redirecting it to *Saa3* (Serum Amyloid A3) and *Acod1* (Aconitate Decarboxylase 1).^29^ Therefore, we next measured transcript levels of *Nfatc1, Saa3, and Acod1* in OCs from both WT and CD14KO after treatment with LPS. Treatment of WT OCs with LPS led to increased expression of *Saa3* (100-fold, p<0.0001) and *Acod1* (4-fold, p<0.0001), as well as decreased expression of *Nfatc1* (70% reduction, p<0.0001). CD14-deficient OCs showed no changes in gene expression with LPS treatment (**Fig. 5E**), and notably *Nfatc1* transcription was completely preserved. Together these results suggest that, in response to a low level LPS/TLR4 challenge, RANKL-mediated OC differentiation is inhibited primarily through the MyD88-pathway, and CD14-deficiency protects against this inhibition by maintaining *Nfatc1* expression.

## Discussion

CD14-deficient mice are partially protected from post-injury SCB remodeling in the DMM model.^15^ Given the relationship between SCB pathology, pain and progression in OA, we sought to better understand the mechanism behind this protection. Here, CD14-deficient mice demonstrated delayed SCB remodeling at 2 weeks post-DMM, reflected in delayed post-injury increases in bone volume, bone mineral density, and trabecular thickness. We further observed that CD14-deficient mice had increased SCB osteoclast content and altered osteoclast/osteoblast ratios 2 weeks post-DMM. This suggests that the delayed bone remodeling could be due to an increase in osteoclast presence in the CD14-deficient subchondral bone, combating the post-DMM increases in bone volume commonly seen in this model.^21^ Additionally, treating WT mice post-DMM with an anti-CD14 mAb revealed lasting protection against pathologic subchondral bone remodeling in the joint, pointing toward the translational potential of CD14 as a therapeutic target. *In vitro*, we observed that CD14-deficient osteoclasts differentiated earlier and more robustly than WT osteoclasts in response to RANKL, but died earlier as well, indicating a similar osteoclastic lifespan between genotypes. In addition, cells from both strains possessed similar cumulative resorptive function, suggesting that the effect of CD14 loss was largely on the rate of differentiation. Bulk RNA sequencing revealed a signature of decreased IFN-I signaling in CD14-deficient OCs. Given the reported inhibitory role IFN-I has on OC differentiation, we used a neutralizing anti-IFNAR1 antibody to test whether effects of CD14-deficiency were mediated by IFN-I. This treatment rescued OC differentiation in WT cells to the levels of CD14-deficient cells, and OCs from IFNAR1KO and IRF3/7KO strains phenocopied the accelerated osteoclastogenesis seen in the setting of CD14-deficiency. Taken together, these findings demonstrate that in response to RANKL, CD14-deficient osteoclasts differentiate earlier due to decreased IFN-I signaling. It has been previously described that, upon RANKL stimulation, osteoclasts release IFN-I as an autoregulatory mechanism.^30^ Our work suggests that the production of IFN-I after RANKL stimulation is dependent upon TLR/CD14 signaling, potentially resulting from endogenous TLR ligands being produced by stressed, differentiating cells. Endotoxin quantification of culture media revealed a very low LPS concentration of 1 pg/mL, unlikely to have an effect in our studies. Further, we show that differentiating osteoclasts produce S100A8/9 as well as Peroxiredoxin-1, both known TLR ligands that could play a role in this autoregulatory mechanism (**Fig. S5)**.^29,31^ Whether this TLR-dependent, autoregulatory phenomenon plays a role in OC differentiation *in vivo* remains to be explored.

CD14 is a co-factor for TLR4 that sensitizes the receptor to respond to low concentrations of ligands, and one such ligand present in OA synovial fluid at low concentrations is the TLR4 ligand LPS.^10^ To test whether CD14 deficiency reduces effects of LPS on OC, we exposed cells from both strains to LPS during differentiation. WT OC differentiation was inhibited with LPS stimulation, as reported previously, and a TLR4 inhibitor reversed this inhibition.^17^ However, CD14-deficient OC differentiation was not inhibited in the presence of LPS, indicating that it is a required co-factor mediating the inhibitory effects of low levels of LPS. Given our results in the absence of LPS stimulation, we again used the anti-IFNAR1 antibody to probe the role of IFN-I in LPS-mediated inhibition of osteoclastogenesis. However, we found no effects of IFNAR1 blockade in the presence of LPS. Further, the IFNAR1KO and IRF3/7KO OCPs did not phenocopy the CD14-deficient OCPs after TLR4 stimulation, as they were still sensitive to the inhibitory effects of LPS. This suggests that, in the presence of LPS, the protective effects of CD14-deficiency on LPS-mediated OC inhibition is not dependent on IFN-I.

LPS binding to TLR4 results in activation of two main signaling pathways. Signaling through TLR4 can occur through MyD88-dependent and -independent pathways, depending on the receptor and receptor complex engaged. Signaling through the MyD88-dependent pathway in OCPs leads to activation of NF-KB signaling and the inhibition of NFATc1, the master regulator of osteoclastogenesis, by diversion of NF-κB away from *Nfatc1* and redirecting it to *Saa3* (Serum Amyloid A3) and *Acod1* (Aconitate Decarboxylase 1).^29^ Activation of the MyD88-independent pathway leads to some degree of NF-κB activation, but primarily acts through IRF3/7 signaling and the production of Type I Interferons (IFN-I), which subsequently inhibits c-Fos, a co-factor for *Nfatc1*.^32^ At low LPS concentrations, CD14 is necessary for signaling through the MyD88-dependent pathway. Conversely, CD14 is necessary regardless of the concentration of ligand for MyD88-independent signaling.^14^ Therefore, it is possible that CD14-deficiency may decrease TLR4 signaling in response to LPS through both the MyD88-dependent and the MyD88-independent (IFN-I producing) pathway. In this study, the decreased expression of *Saa3* and *Acod1*, and preservation of *Nfatc1* transcription, in CD14-deficient osteoclasts exposed to LPS, is consistent with reduced signaling through the MyD88-dependent pathway. Taken together with our findings showing no effects of IRF3/7KO or IFNAR1KO and neutralization, this suggests that the protection from LPS inhibition in the CD14-deficient OCPs is mediated primarily through decreased MyD88-dependent signaling.

There are several limitations of the current study to point out. First, our data suggests, but does not definitively demonstrate, that the MyD88-dependent pathway is solely responsible for the differential effects of LPS on WT and CD14KO cells, as both TRIFKO and MyD88KO cells showed profound suppression of RANKL-mediated osteoclastogenesis at baseline and did not expand sufficiently to test this hypothesis. Published work shows the necessity of MyD88 in OC differentiation when in co-culture with OB.^33^ It is possible that some activation of this pathway is necessary to initiate differentiation, and future work can utilize additional methods to probe the differential effects of this pathway on OC differentiation in the absence and presence of LPS. Further, LPS is only one OA-relevant TLR4 ligand, and it cannot be assumed that other OA-relevant TLR PAMPs and DAMPs will have the same effects on OC differentiation.^9^ Future work will shed light on the effects of other OA-relevant TLR ligands on WT and CD14-deficient OC differentiation. Lastly, more work is required to confirm the role of MyD88-dependent and - independent signaling in OA SCB sclerosis *in vivo*. Still, the present work sheds light for the first time on an important role for IFN-signaling downstream of CD14 in mediating pathologically reduced osteoclastogenesis after joint injury. This opens up new potential pathways to therapeutically target aberrant bone remodeling in the setting of joint injury and PTOA.

## Acknowledgements

We thank Implicit Bioscience Ltd (Brisbane, Australia) for supplying the murine anti-CD14 antibody (clone biG53) used in the DMM experiments, which was obtained under an MTA. We thank Saeed Jerban, PhD for the custom MATLAB code used to analyze the femoral subchondral bone. We additionally thank Jonathan Schug, John Tobias, and the research staff at the Penn Genomics and Sequencing Core. Lastly, we’d like to thank Sam Chauvin for transporting the IFNAR1KO and IRF37DKO mice.

