## Supplemental Materials for "CD14-deficiency protects against osteoarthritic subchondral bone sclerosis via enhanced osteoclastogenesis following joint injury"

**Supplemental Methods**

**LPS assay:** To assay for LPS contamination in culture media, the Pierce Chromogenic Endotoxin Quant Kit (ThermoFisher) was used.

**S100A8/9 ELISA:** A commercially available ELISA (ThermoFisher) was used to measure S100A8/9 levels in supernatants from osteoclast cultures on days 2 and 4 post-RANKL addition, per the manufacturer’s instructions.

**Supplemental Results**

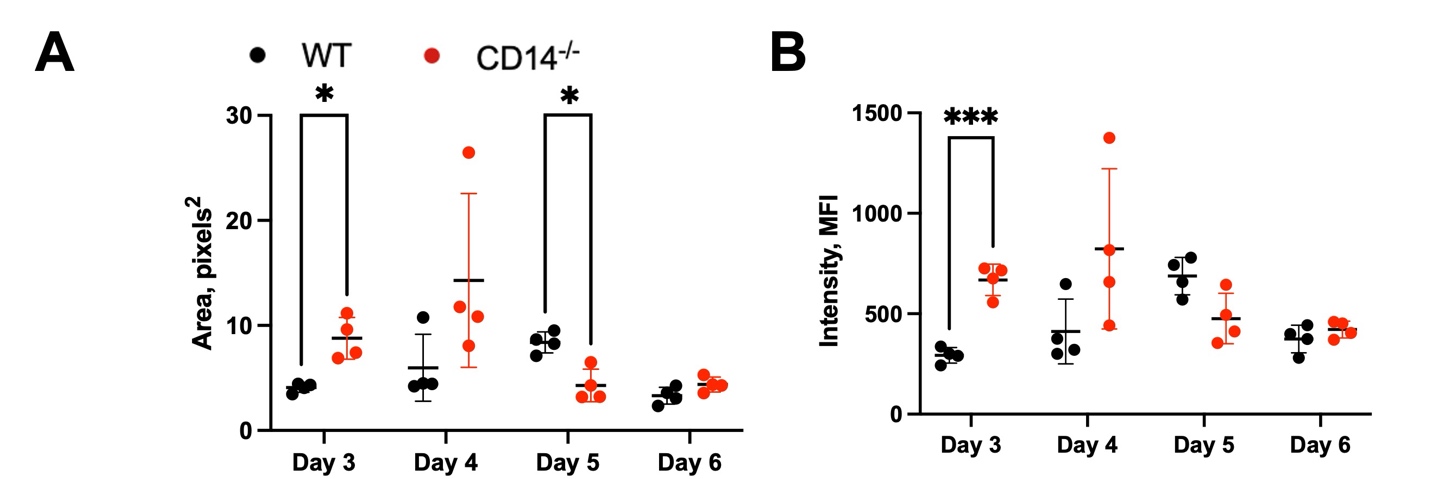

**Figure S1: WT and CD14-deficient osteoclasts increase in size and DAPI intensity as they differentiate.** Mean osteoclast area (A) and DAPI intensity (B) of WT and CD14-deficient osteoclasts (B). *p<.05, ***p< .001

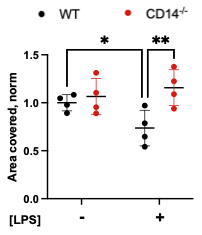

**Figure S2: LPS-induced inhibition on WT osteoclast differentiation is partially rescued with TLR4 inhibition.** Percent area of WT and CD14KO osteoclasts with and without LPS on day 4 post-RANKL stimulation, treated with CLI-095. *p<0.05, **p<0.01

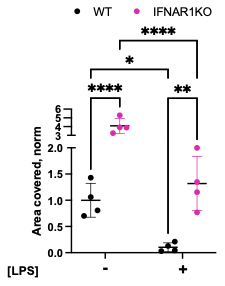

**Figure S3: LPS significantly inhibits IFNAR1KO osteoclast differentiation.** Percent area of WT and IFNAR1KO osteoclasts with and without LPS on day 3 post-RANKL stimulation. *p<0.05, **p<0.01, ****p<0.0001

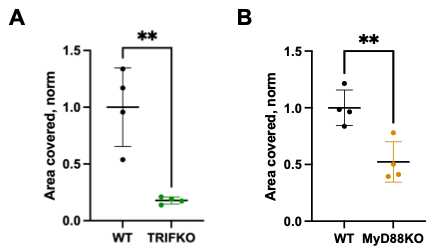

**Figure S4: OCPs isolated from TRIFKO and MyD88KO mice show reduced OC differentiation.** Percent area of osteoclasts on day 3 after RANKL addition, for OCPs derived from WT, TRIFKO (A), and MyD88KO (B). **p<0.01

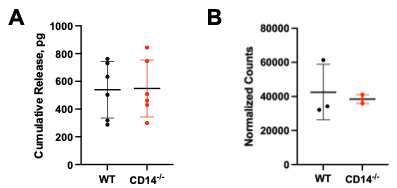

**Figure S5: Potential TLR ligands S100A8/9 and Peroxiredoxin-1 are produced by WT and CD14-deficient osteoclasts.** Cumulative release of S100A8/9 into media by WT and CD14-deficient osteoclasts on day 4 following RANKL stimulation (A). Transcript counts of peroxiredoxin-1 in WT and CD14-deficient osteoclasts, taken from bulk RNA sequencing (B).
